# Position Effects Influencing Intrachromosomal Repair of a Double-Strand Break in Budding Yeast

**DOI:** 10.1101/114850

**Authors:** Ruoxi W. Wang, Cheng-Sheng Lee, James E. Haber

## Abstract

Repair of a double-strand break (DSB) by an ectopic homologous donor sequence is subject to the three-dimensional arrangement of chromosomes in the nucleus of haploid budding yeast. The data for interchromosomal recombination suggest that searching for homology is accomplished by a random collision process, strongly influenced by the contact probability of the donor and recipient sequences. Here we explore how recombination occurs on the same chromosome and whether there are additional constraints imposed on repair. Specifically, we examined how intrachromosomal repair is affected by the location of the donor sequence along the 812-kb chromosome 2 (Chr2), with a site-specific DSB created on the right arm (position 625kb). Repair correlates well with contact frequencies determined by chromosome conformation capture-based studies (*r* = 0.85). Moreover, there is a profound constraint imposed by the anchoring of the centromere (*CEN2,* position 238kb) to the spindle pole body. Sequences at the same distance on either side of *CEN2* are equivalently constrained in recombining with a DSB located more distally on one arm, suggesting that sequences on the opposite arm from the DSB are not otherwise constrained in their interaction with the DSB. The centromere constraint can be partially relieved by inducing transcription through the centromere to inactivate *CEN2* tethering. In diploid cells, repair of a DSB via its allelic donor is strongly influenced by the presence and the position of an ectopic intrachromosomal donor.

**Author Summary:** A double-strand break (DSB) on a chromosome can be repaired by recombining with an ectopic homologous donor sequence. Interchromosomal ectopic recombination is strongly influenced by the three-dimensional arrangement of chromosomes in the nucleus of haploid budding yeast, that is strongly influenced by the probability of chemical cross-linking of the donor and recipient sequences. Here we explore how recombination occurs on the same chromosome. We examined how intrachromosomal repair is affected by the location of the donor sequence along the 812-kb chromosome 2 (Chr2), with a site-specific DSB created on the right arm (position 625kb). Repair correlates well with contact frequencies determined by chromosome conformation capture-based studies (*r* = 0.85). Moreover, there is a profound constraint imposed by the anchoring of the centromere (*CEN2*, position 238kb) to the spindle pole body. Sequences at the same distance on either side of *CEN2* are equivalently accessible in recombining with a DSB located more distally on one arm, suggesting that sequences on the opposite arm from the DSB are not otherwise constrained in their interaction with the DSB. The centromere constraint can be partially relieved by inducing transcription through the centromere to inactivate *CEN2* tethering. In diploid cells, repair of a DSB via its allelic donor is strongly influenced by the presence and the position of an ectopic intrachromosomal donor.

## Introduction

A fundamentally important step in the repair of a broken chromosome by homologous recombination is the identification and use of a homologous donor sequence to repair the DSB [Agmon, 2013 #12545;Lee, 2016 #12798;Inbar, 1999 #10663;Haber, 2013 #11987;Coic, 2011 #10624;Lichten, 1989 #643](1–6). In eukaryotes, DSBs are processed by exonucleases to expose 3’-ended single-stranded regions upon which Rad51 recombination protein is loaded and forms a nucleoprotein filament. The Rad51 filament, like its bacterial RecA counterpart, then interrogates other sequences in the genome to locate a homologous segment with which it can promote strand invasion to form a displacement loop and then initiate DNA synthesis using the homologous sequence as a template to repair the DSB. How the search for homology – on a sister chromatid, a homologous chromosome or at an ectopic site –is accomplished remains a subject of active investigation. Several lines of evidence suggest that the search is more efficient intrachromosomally – at least over modest distances of 100- 200 kb [Lichten, 1989 #643;Lee, 2016 #12798](2,6), but this question needs to be explored in more detail.

Recently we developed an ectopic donor assay to study DSB repair efficiency in haploid *Saccharomyces cerevisiae* [Lee, 2016 #12798](2). A site-specific DSB, created by the galactose-inducible HO endonuclease, could be repaired by a single ectopic donor sequence, which shares 1 kb homology with either side of the break and is located elsewhere in the genome. By placing a donor at 20 different locations throughout the genome, we showed that the efficiency of interchromosomal recombination was strongly correlated with the likelihood that the donor region would come into contact with the recipient site, based on chromosome conformation capture analysis [Duan, 2010 #10805;Lee, 2016 #12798](2,7). In some cases, a donor site that was quite inefficient when used to repair the DSB on Chr5 became much more efficient when confronted with a different DSB induced on Chr2, consistent with the differences in its contact frequencies with the regions surrounding the two break sites. Studies by Agmon et al. [Agmon, 2013 #12545](1) and by Zimmer et al. [Zimmer, 2011 #12012] (8) also reached similar conclusions, with focus on the recombination when both DSB and donor are located at pericentrimeric or subtelomeric regions.

Here we have extended our analysis to examine the correlation between contact frequencies and repair for intrachromosomal events. We find again a strong correlation with contact frequency but also see additional constraints imposed by the centromere and by the very high level of contacts made by nearby intrachromosomal sequences.

## Results

### Intrachromosmal GC is subjected to chromosome organization

In our previous study, we mainly focused on the correlation between chromosome organization and recombination frequencies in interchromosomal noncrossover gene conversion events [Lee, 2016 #12798] (2). A DSB was induced within *leu2* sequences inserted on Chr5, while a homologous 2-kb *LEU2* donor was placed at 4 positions on the same chromosome or at 20 locations on different chromosomes. Repair occurs predominantly by synthesis-dependent strand annealing in which by a patch of DNA newly copied from the donor to replaces the 117-bp HO cleavage site sequences [Lee, 2016 #12798][Ira, 2006 #2857] (2,9). We and others have observed that intrachromosomal gene conversion occurred generally more efficiently and with a faster kinetics than interchromosomal events [Lee, 2016 #12798;Agmon, 2013 #12545][Mehta, 2017 #13041] (1,2,10).

To examine intrachromosomal repair in more detail, we constructed a series of 12 strains, in which a DSB was created within a 2-kb *LEU2* gene inserted 625kb from the left end on chromosome 2 (Chr2) and a donor was inserted at different sites across the chromosome (Figure 1A). The DSB, created by galactose-inducible expression of the HO endonuclease gene, is situated 387 kb from *CEN2* and 187 kb from the right telomere (http://www.yeastgenome.org/). The efficiency of DSB repair of each of the 12 strains was measured by plating cells on YEP-galactose plates to continuously induce the HO endonuclease, compared to the same number of cells plated on glucose-containing medium. Virtually all of the survivors repaired the DSB by ectopic gene conversion rather than by nonhomologous end-joining, which occurs only in 0.2% of cases [Lee, 2016 #12798](2). Cell viabilities among the 12 strains ranged from 9% to 89%; because these repair events occur on the first cell cycle, the observed frequencies are equivalent to rates.

**Figure 1:**
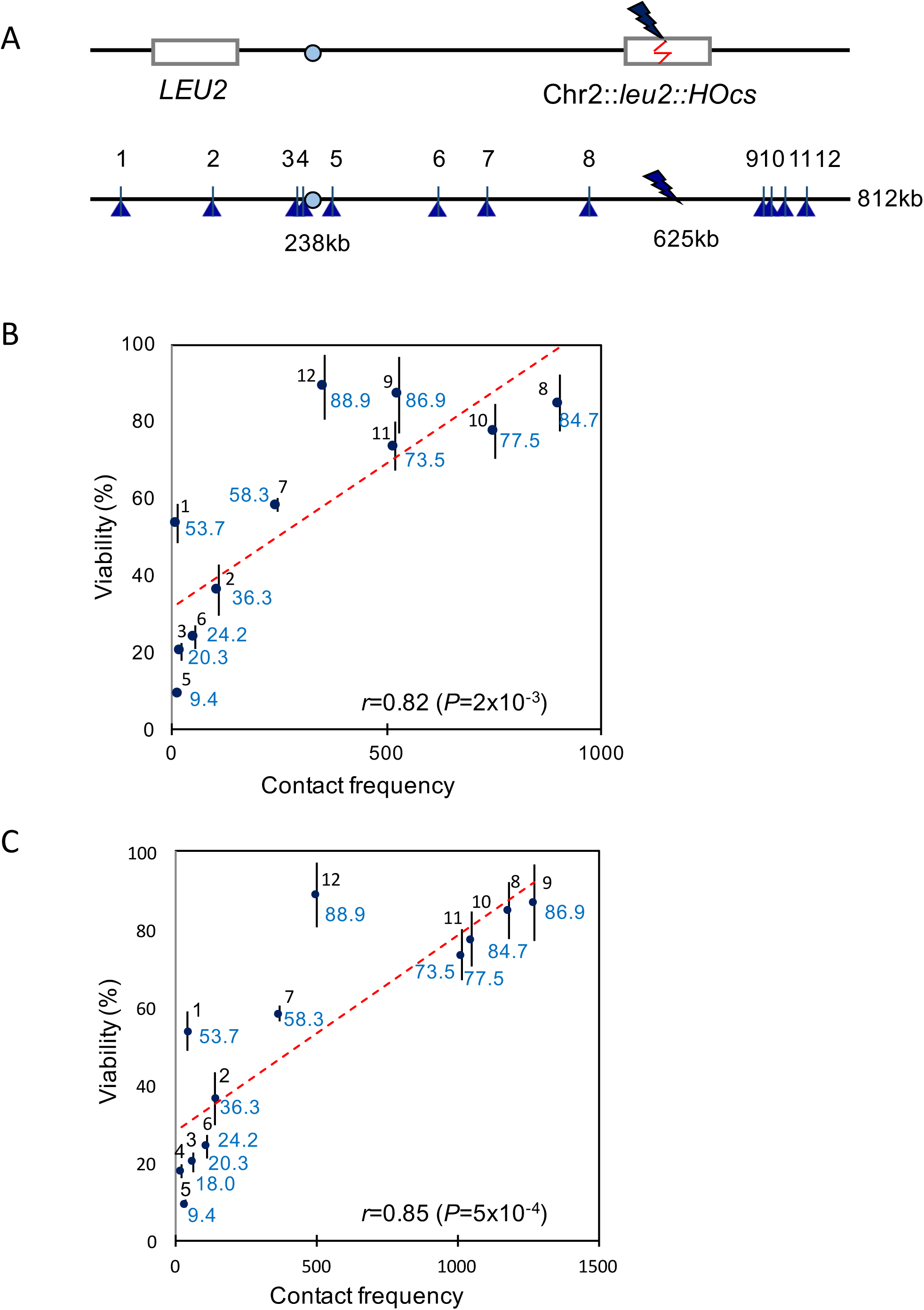
Viability assay to assess repair efficiency for 12 intrachromosomal loci. (A) The scheme of viability assay. The *leu2::HOcs* was inserted at 625 kb on Chr2. The DSB could be repaired by an ectopic *LEU2* donor inserted on the same chromosome. The locations for the 12 donors were shown along Chr2. (B and C) Correlation between cell viability (%, shown in blue) and total contact frequency using ±25kb window size around Chr2-DSB and ±10 kb(B) or ±20 kb window size around donor(C). Only 11 loci were analyzed in (B) since no productive contact was detected between ±25kb around DSB and ±10 kb around site 4. Error bars indicate one SD from three independent experiments.

Repair efficiencies were then plotted with respect to the total contact frequencies between the DSB region and the donor region, which is calculated by adding up all the interaction reads, measured by [Duan, 2010 #10805] (7), between +/−25kb region surrounding the DSB site and either a +/−10kb (Figure 1B) or +/−20kb (Figure 1C) region surrounding a donor site. Cell viability displayed a strong correlation with the total contact frequency; however, the effect of contact frequency on cell viability approached saturation when donor was within about 100 kb of the DSB, where contact frequencies also reached a maximum (Figure S1B). The calculated correlation coefficient using Pearson correlation analysis was *r* = 0.82 (*P* = 2 × 10^−3^) with +/−10kb window around the donor and *r* = 0.85 (*P* = 5 × 10^−4^) with +/−20kb window around the donor. Thus, intrachromosomal recombination is strongly constrained by the likelihood that two sequences will come into contact, as we saw for interchromosomal events.

We note that when donors were located within 50 kb from a telomere (locus 1 and 12), their measured viabilities were higher than expected based on their contact frequency (Figure 1B and 1C). It has been reported that chromosomal conformation capture methods tend to underestimate productive contacts in subtelomeric regions [Duan, 2010 #10805] (7). Although sequences more than 20 kb from a telomeric anchor appear to be unconstrained [Avsaroglu, 2014 #12111] (11), it seems possible that the underestimation of contact frequencies may explain the higher-than-predicted recombination efficiencies for these two loci. The results for donors placed within 100 kb of the DSB target appear to reflect a plateau, consistent with the leveling off of contact frequencies (Figure S1B).

### DSB repair is constrained by centromere tethering

If one plots the correlation between cell viability and the distance of a homologous *LEU2* donor from the left end of Chr2, it becomes evident that donors located close to the centromere display a low repair rate compared to donors located within a chromosome arm (Figure 2A). Indeed, the efficiencies of repair are in agreement with the idea that the two chromosome arms are in the Rabl orientation [Jin, 2000 #137] (12), with the centromere anchored at the spindle pole body (SPB) (Figure S1A). In budding yeast the centromere remains attached to the SPB throughout the cell cycle [Tanaka, 2007 #147] (13). Interestingly, if one re-plots recombination efficiencies as a function of the distance from *CEN2,* it becomes apparent that the left arm - where the homologous *LEU2* locations are significantly more linearly distant from the DSB itself – behaves as if these sites are as accessible as those on the right arm (Figure 2B). These results suggest that the tethering of the telomere of the left arm does not prevent sequences from interacting with the DSB on the opposite arm as efficiently as sites on the right arm, when the sites are equally distant from the centromere.

**Figure 2:**
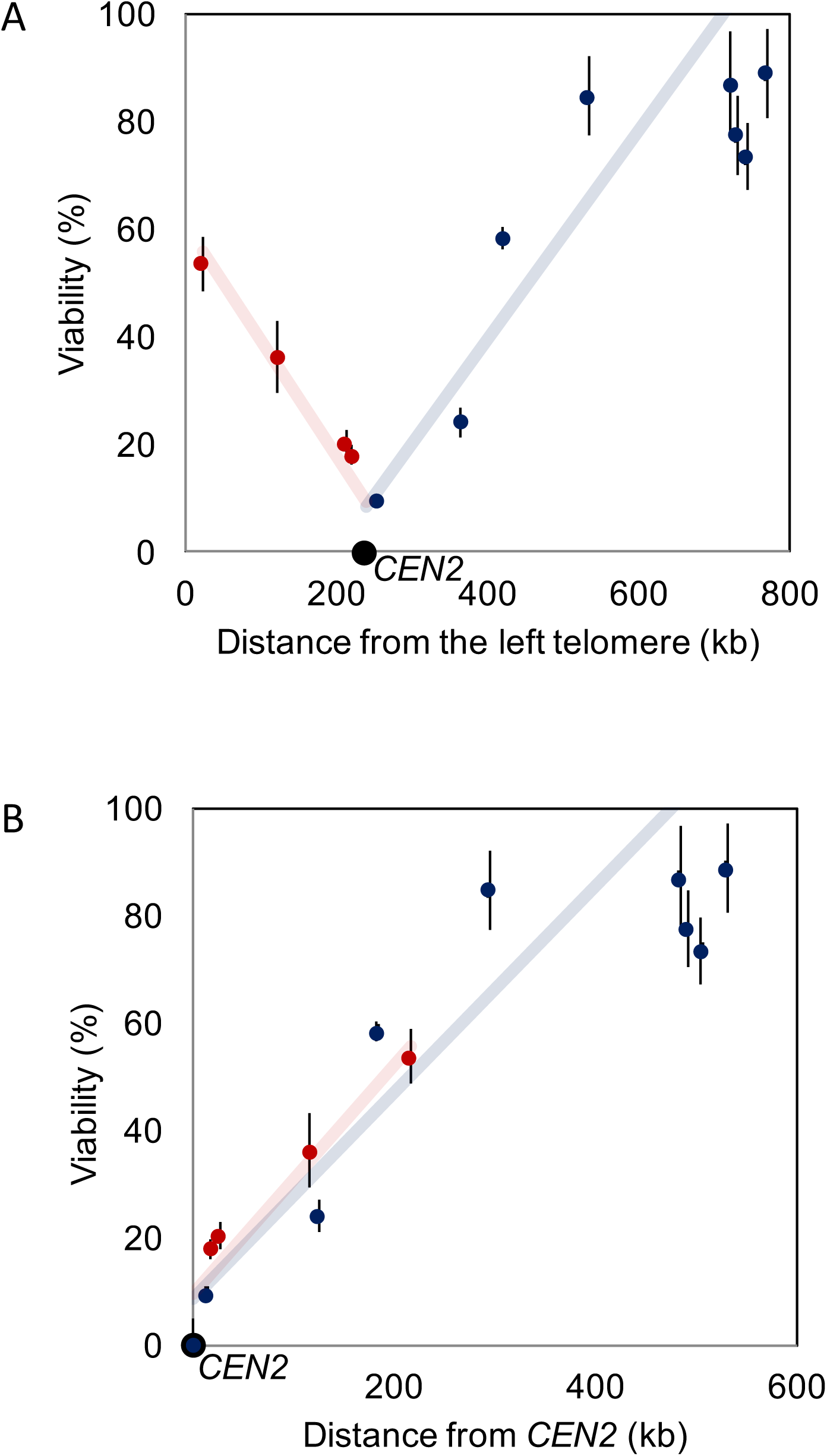
Correlation between cell viability and distance of a homologous *LEU2* donor from (A) the left telomere and (B) the centromere *(CEN2).* Pearson's correlation test were conducted for either side of *CEN2* (including *CEN2)* respectively. Donor sites 1012 (729kb, 742kb and 768kb) are excluded from the analysis because viability had reached a plateau. *r* = 0.93 for right side of *CEN2,* and *r* = 0.95 for left side of *CEN2.*

To further explore if the correlation between genomic distance and cell viability is affected by centromere tethering, we enquired whether detaching the centromere from SPB would alter the pattern of repair we observed in wild type strains. Cohesin is an essential protein complex that facilitates spindle attachment to the centromere. Mcm21 is a non-essential kinetochore component of the COMA complex [Ortiz, 1999 #13034] (14) that is responsible for the enrichment of cohesin at the pericentromeric region. Deletion of *MCM21* results in a partial dispersal of kinetochores from the normal cluster around the SPB, but does not prevent relatively normal chromosome segregation [Tsabar, 2016 #12794] (15). However, deleting *MCM21* did not result in a change in repair efficiency (and thus cell viability) among four of the *MCM21* deletion strains whose donors were close to *CEN2* (Figure 3A), suggesting that the depletion of Mcm21 protein might not be sufficient to fully inactivate the attachment of centromere to the SPB.

**Figure 3:**
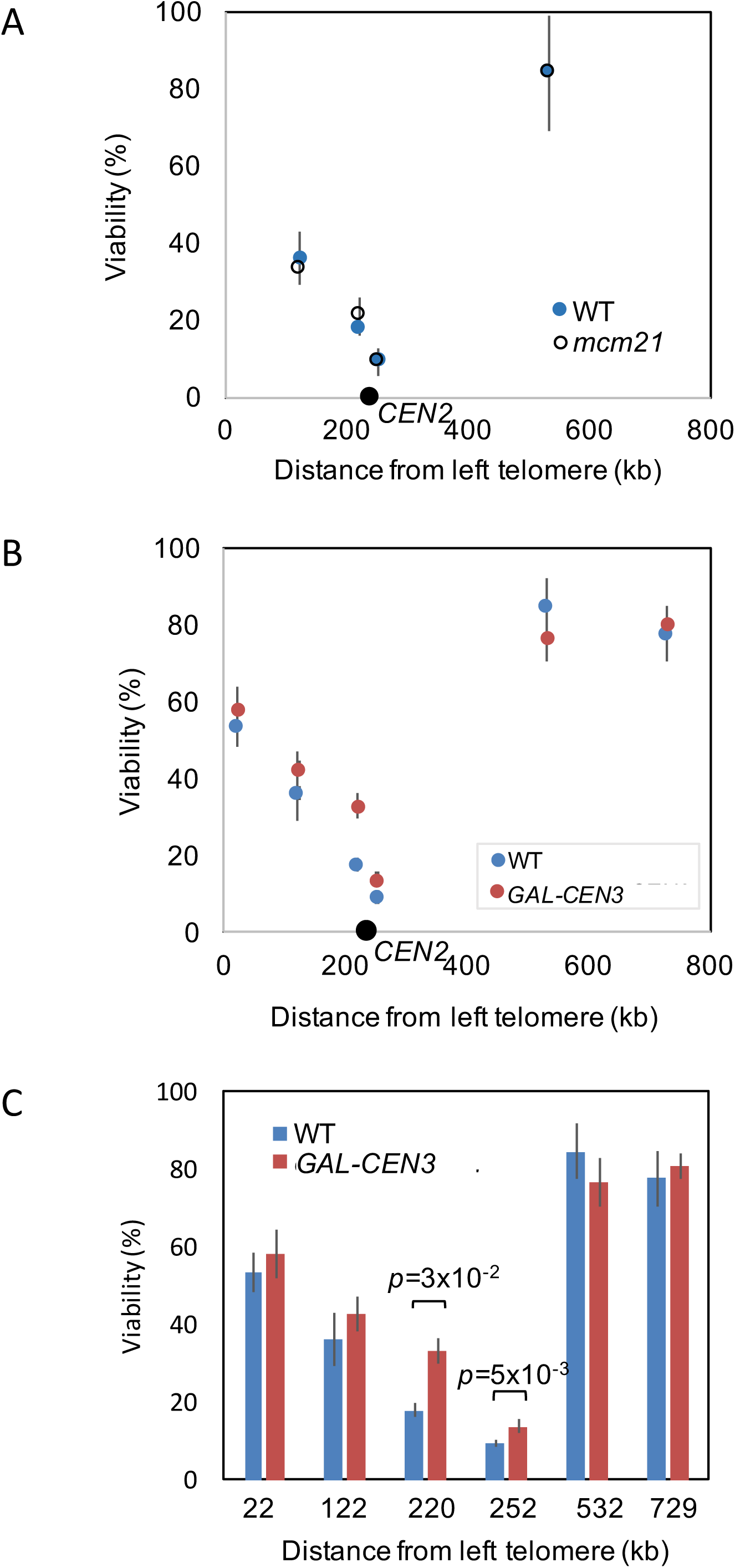
(A) Effect of *mcm21*Δ*A* on viability. (B and C) Effect of *cen2*Δ*A::GAL-CEN3* on viability. Inactivation of *CEN2* significantly increased viability of two donors located close to *CEN2.* Error bars indicate one SD from three independent experiments.

As an alternative way of disrupting kinetochore attachment to the SPB, we introduced a conditionally functional centromere by placing a galactose-inducible promoter upstream of the centromeric DNA sequence [Hill, 1989 #661;Tsabar, 2016 #12794] (15,16). A *GAL::CEN* centromere is functional when cells are grown on glucose-containing plate but its function is impaired when cells are transferred to galactose-containing plate, as the strong transcription disrupts normal assembly of the kinetochore at this centromere. Our recent study of the *GAL-CEN3* construct showed that sister chromosomes properly segregated only 1/3 of the time, and then only after some delay [Tsabar, 2016 #12794] (15). We therefore replaced *CEN2* with *cen2::GAL-CEN3* in several of the intra-chromosomal donor strains (Figure 3B). Placing cells on galactose, which simultaneously induced HO endonuclease expression and inactivated the Chr2 centromere, had no significant effect on donors located far from *CEN2,* but did significantly raise the level of repair of two loci near *CEN2* (Figure 3C).

To confirm that transcription did indeed perturb centromere function in these strains we carried out pedigree analysis to measure missegregation of the chromosome by the appearance of daughter cells that failed to inherit Chr2 [Tsabar, 2016 #12794] (15). To be sure that segregation was not also influenced by the HO cleavage, we modified strains by removing the HO cleavage site, so that only the *GAL::CEN* would be affected by addition of galactose. This was accomplished by transforming the strain with a HPH-marked plasmid expressing both Cas9 and a guide RNA targeted to the HO cleavage site. Transformants grown on glucose medium proved to have lost the cleavage site, by DSB-induced gene conversion using the ectopic *LEU2* donor (data not shown). For strains with wild type *CEN2* (cutsite-deleted derivatives of strains YWW113 and 119), both mother and daughter cells gave rise to colonies in each of 45 cases. For the modified strain YWW216 (donor at 220 kb, 18 kb from *CEN2*), lacking the HO cutsite, 16 of 29 daughters failed to produce colonies, and for modified strain YWW231 (donor at 252 kb, 14 kb from *CEN2*), 7 out of 20 daughter cells failed to give rise to colonies. Finding approximately 1/3 of pedigrees failing to properly disjoin the *GAL::CEN* chromosome is consistent with our previous study and confirms that when galactose was added to induce HO it also would disrupt normal *CEN* function [Tsabar, 2016 #12794] (15).

### Interchromosomal versus intrachromosomal repair of a DSB

In diploid yeast, mitotic homologous chromosomes are not evidently paired with each other although some preferential interactions have been reported [Burgess, 1999 #1589;Burgess, 1999 #1590;Lorenz, 2003 #13038] (17–19). We created diploids to ask how the presence of a competing allelic donor would affect repair via an intrachromosomal site. Whereas the ectopic intrachromosomal donor shares only 1 kb on each side of the DSB, the homologous chromosome shares the entire chromosome arm. The diploid strains were constructed by mating strains that carried the *leu2::HOcs* at 625kb and an intrachromosomal ectopic *LEU2* donor at different locations with a strain carrying a *URA3* selectable marker and a *leu2-K* donor at 625 kb; that is, at the allelic position to the DSB (Figure 4A). Normal *MAT* sequences were deleted (see Materials and Methods). In these strains, viability was nearly 100% as expected for a diploid where an unrepaired DSB and chromosome loss would still lead to a viable aneuploidy [Malkova, 1996 #333] (20). We assessed the use of the intrachromosomal ectopic *(LEU2)* and allelic *(leu2-K)* donors by PCR-amplifying the repaired locus followed by *Kpn*I digestion, both in pools of cells (Figure S2) and from colonies of individual recombinants (Figure 4B). HO cleavage is nearly 100% efficient so no amplification occurs from the unrepaired *leu2::HOcs* site.

**Figure 4:**
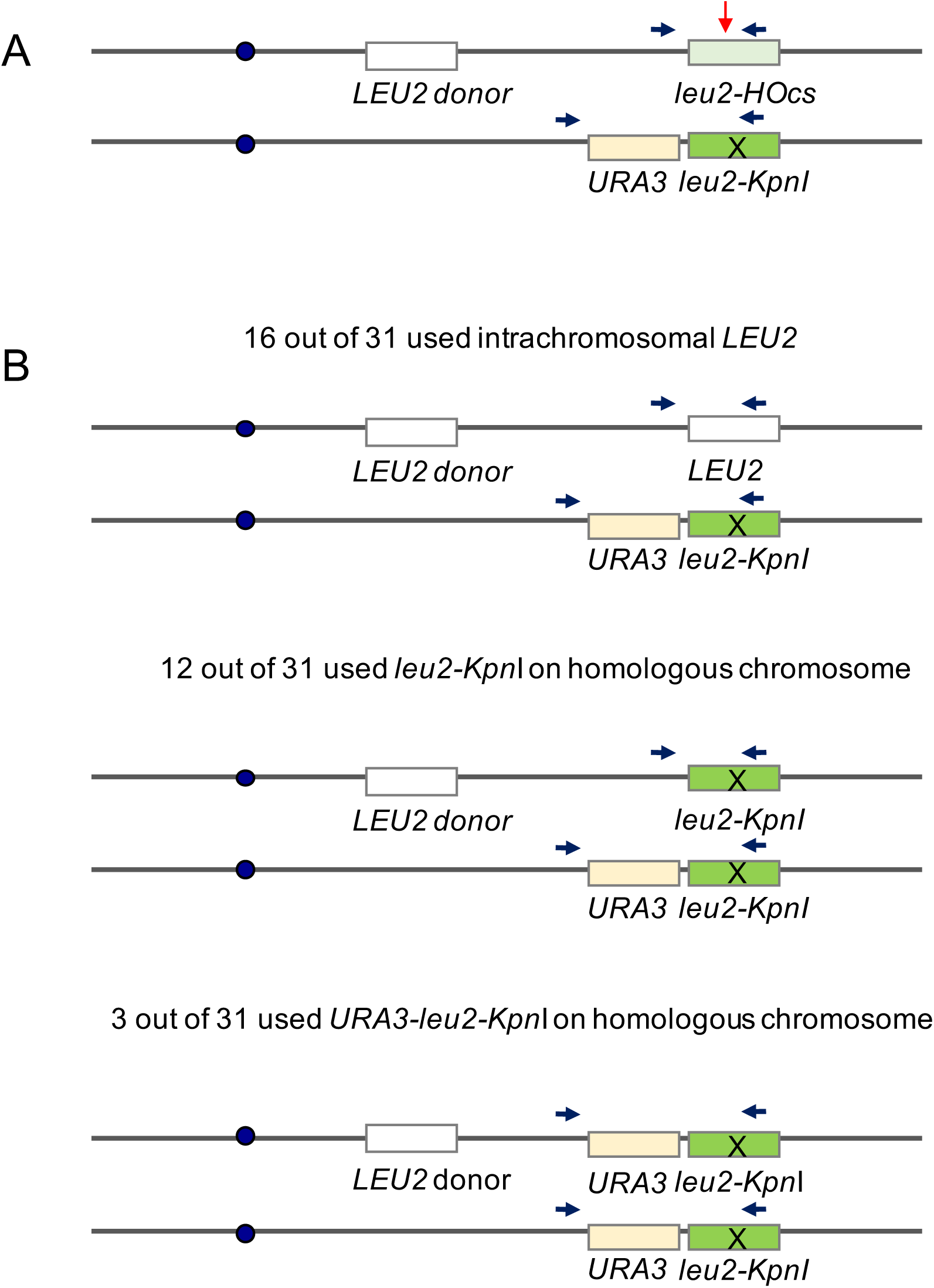
(A) Scheme for DSB repair in diploid strains. The DSB could be repaired by gene conversion using an ectopic intrachromosomal *LEU2* sequence, an allelic *leu2-*Kpnl sequence, or the homologous sequence outside of the leu2-Kpni. (B) Usage of ectopic and allelic donors assessed from 31 colonies of individual recombinants.

The use of the ectopic (KpnI+) donor was substantially reduced in each of three examples (Figure S2C), as compared to the use measured in a haploid strain by a viability assay (Figure 1B). In pooled cells, for example, for site 2 (122kb), which was 36.3% viable in the haploid strain assay, only 12.3% of the repair events used this intrachromosomal donor. For site 8 (532kb), which was a highly efficient donor (84.7%) in the haploid assay, its use was decreased to 57.7% in the diploid, with the remainder coming from the allelic locus. The PCR-*KpnI* assay used in Figure S2 slightly underestimates the use of the allelic donor because it fails to capture the fraction of allelic gene conversion events that co-convert *leu2-K* and the adjacent *URA3* marker (as illustrated in Figure 4B, bottom panel). This larger insertion was not amplified under the PCR conditions used to assay the population of recombinants, as shown in Figure S2B. To assess the proportion of the three types of possible outcomes, we analyzed individual repair events (Figure 4B), where we used PCR conditions that recover all of the relevant products. Of 31 events, 16 (51.6%) used the intrachromosomal donor, while 12 repaired the DSB without co-converting the adjacent *URA3* marker and 3 co-converted *URA3* along with *leu2-K.*

The ectopic donor shares 1 kb homology on either side of the DSB whereas the allelic donor has extensive homology on both sides of the break (with a 1-kb insertion on one side). Previously we have shown that increasing homology from 1 kb to 2 or 3 kb on each side of the DSB had a very significant effect on the efficiency of ectopic DSB repair [Lee, 2016 #12798] (2); the data here are consistent with the idea that sharing more extensive homology, even if interrupted on one side by a heterology, has a highly significant effect on the likelihood that a donor will be successful in repairing the DSB; but the intrachromosomal site remained the preferred donor.

## Discussion

In haploid yeast genome, the sixteen chromosomes adopt a preferential 3D conformation with centromeres clustered at spindle pole body and telomeres loosely associated at the nuclear envelope, the so-called Rabl configuration [Taddei, 2010 #13040] (21). Our previous work and that of others have shown that 3D nuclear architecture is a key factor that influences the rate and efficiency of interchromosomal DSB repair, with a striking correlation between repair and the estimation of the physical distance of two DNA fragments in the genome (contact frequency). These studies also demonstrated that a site that served as an efficient donor to repair a DSB at one location could be a very inefficient donor when the site of DSB (with the same homologous sequences) was moved to a different chromosome; thus most donor sites were not “hot” or “cold” because of local chromatin features.

Here we show that chromosome conformation capture data also provide strong predictions for intrachromosomal DSB repair frequencies. Sites within approximately 100 kb of the DSB, which show very high levels of contact frequency, reach a plateau in their ability to recombine, but beyond that distance repair roughly diminishes with distance as the donors are placed closer to the centromere. However, at increasing distance from the centromere on the opposite chromosome arm, repair frequencies increase. This pattern is consistent with the Rabl configuration of chromosomes and, further, that a site 200 kb on the left arm is approximately as able to recombine as one on the right arm, despite being much further away as viewed along the chromosome itself. These results suggest that the left arm is not tethered away from the DSB site. The high level of accessibility of sites on the left arm that are distant from the centromere is not evident in the Hi-C data, which is swamped by interactions close to the site of interest (Figure S1B).

Our results demonstrate a strong constraint on the ability of centromere-proximal sequences to recombine with distant loci, although Agmon et al. [Agmon, 2013 #12545] (1) showed that recombination between centromere-adjacent sequences on different chromosomes is efficient, consistent with the bundling of centromere-adjacent sequences held by the cluster of centromeres at the SPB. Our results contrast with those of Agmon et al., who suggested that recombination involving one interstitial element should not be constrained by any tethering effects.

Despite reports that deletion of *MCM21* leads to the partial dislocation of centromeres from the SPB [Ortiz, 1999 #13034] (14), this deletion did not relieve the constraint of centromere proximity in DSB repair. However, disruption of *CEN2* function by galactose-induced transcription proved to cause a modest but statistically significant increase in the ability of centromere-proximal sequences to recombine. We note that even with *GAL::CEN* disruption, 2/3 of cells are able to maintain proper chromosome segregation, so the effect of disrupting *GAL::CEN* would not necessarily be expected to have a larger consequence.

Previously we had shown that there could be a strong completion between an intrachromosomal donor and a competitor at an allelic site for spontaneous mitotic recombination [Lichten, 1989 #643;Lee, 2016 #12798] (2,6). Here, we show that this conclusion holds true for events known to be initiated by a site-specific DSB, depending on the contact frequency between intrachromosomal sites. It will be interesting to assess these results in more detail when contact probabilities have been determined in diploid strains.

## Materials and Methods

### Strains

All strains were derived from YCSL305 *(ho hmlΔ::ADE1 mataΔ::hisG hmr*Δ*::ADE1 leu2::KAN ade3*::GAL::HO *ade1 lys5 ura3-52 trp1 Chr2.625kb::leu2::HOcs).* The specific locations of the donors inserted on Chr2 and the derivatives containing either *mcm21*Δ*A* or a *GAL::CEN* replacement of *CEN2* are presented in Table S1. A NAT-MX cassette amplified from pJH1513 was inserted at the specific donor location and was then replaced by a TEFp-LEU2-TEFt fragment through homologous recombination. Deletion of *MCM21* was accomplished by transforming cells with a PCR-amplified NAT-marked deletion, copied from the yeast genome knockout collection [Chu, 2008 #153]. The *URA3::GAL::CEN3* sequence was amplified from pJH870 using PCR primers cen2::GAL-CEN3 p4 and cen2::GAL-CEN3 p5 to replace the CEN2 region. The sequences of the primers used in strain construction are presented in Table S2.

Diploid strains were constructed by mating **a**-like strains (deleted for *MAT)* that carried the *leu2::HOcs* at 625kb and an intrachromosomal ectopic *LEU2* donor at different locations with another *MAT*-deleted strain carrying a *URA3* selectable marker and a *leu2-K* donor at 625 kb (that is, at the allelic position to the DSB). Mating was accomplished by transforming the second strain with a *TRP1-marked MATα* plasmid, which was not retained after mating.

### Growth conditions

Single colonies were inoculated into YP-Lactate medium and grew to log phase at 30 ^°^C. Viability assays were carried out as described by Lee et al. [Lee, 2016 #12798]. The viability was calculated as the number of colonies that grew on YEP-galactose medium divided by the number of cells grew on YEPD medium. Three biological replicates were performed on each strain. Pearson's correlation test was conducted between viability and contact frequency.

### Pedigree analysis

The disruption of normal Chr2 segregation in the *cen2::GAL::CEN3* strain was determined by pedigree analysis as previously described. Individual unbudded (G1) cells were micromanipulated and allowed to grow until mother and daughter cells could be separated. The subsequent growth of the mothers and daughters was observed after 24 h. Daughter cells that failed to inherit Chr2 at the first cell division failed to proliferate beyond another cell division, whereas a normal cell or a mother cell that inherited an extra copy of Chr2 grew into a microcolony.

### PCR analysis for diploid strains

Single colonies were inoculated into 4ml of YP-Lactate medium and grew at 30 °C overnight. The culture was diluted and grew to log phase. Then DSB was induced by adding 20% galactose to a final concentration of 2%. The repaired region was amplified from purified genomic DNA using flanking primers Mcm7p3 and Leu2p18B (sequences are presented in Table S2). Short PCR extension time was used to avoid the amplification of *URA3* from the homologous chromosome. The PCR amplicon was digested with KpnI overnight. The digested fragments were separated and visualized on agarose gel. The relative usage of intrachromosomal donors was calculated by dividing the sum of intensity of the *KpnI* fragments by the total intensity of all amplicons. The experiments were repeated for three times in each strain.

## Figure Captions

Figure S1: (A) Rabl configuration of a chromosome in budding yeast. The centromere is tethered to the spindle pole body and the telomeres are clustered at the nuclear envelope. (B) Distribution of intrachromosomal contacts to the DSB site (Chr2, 625kb) using ±25 kb window size around the DSB. The contact frequency between the DSB and the donor is determined by adding up all individual contacts around the donor location. The position of the HO cleavage site is given by a red arrow.

Figure S2: (A) Usage of ectopic and allelic donors assessed from a population of cells. The 3kb *URA3-/eu2-Kpn*I sequence was excluded in the PCR-based analysis by using short amplification times, as indicated by smaller arrowheads. (B) An example of donor usage measurement on agarose gel (YWW210, 57.5% intrachromosomal donor usage). The top band (1045bp) represents *leu2-KpnI* repair product. The lower two bands (732bp and 313bp), digested by *KpnI*, represent *LEU2* repair product. The intrachromosomal donor relative usage (%) was calculated as the intensity of the sum of lower two bands divided by the total intensities of the three bands. (C) Plot of intrachromosomal donor relative usage versus contact frequency (±10kb around donor and ±25 kb around DSB). The intrachromosomal donor locations and their corresponding viabilities (%) in haploid strains are shown in blue. Error bars indicate one SD from three independent experiments.

